# A Data-Analysis Pipeline for High-Throughput Systematic Evolution of Ligands by Exponential Enrichment (HT-SELEX) in the Characterization of Telomeric Proteins

**DOI:** 10.64898/2026.03.06.710105

**Authors:** Jonathan D. Williams, Valerie M. Tesmer, Sagarika Kannoly, Hiroki Shibuya, Jayakrishnan Nandakumar

## Abstract

Telomeres are nucleoprotein structures at the ends of eukaryotic chromosomes that safeguard them from triggering inappropriate DNA damage signaling. POT1, a member of the mammalian shelterin complex, binds single-stranded (ss) telomeric DNA and blocks the activation of the ATR kinase-mediated DNA damage response at telomeres. Yet until recently, it was poorly understood how the double-stranded (ds)-ss telomeric junction was protected from DNA damage response factors. An initial study of the DNA-binding activity of human POT1 (hPOT1) using systematic evolution of ligands by exponential enrichment (SELEX) and subsequent investigation revealed that POT1 contains a binding pocket, known as the POT-hole, that binds the 5’ phosphorylated dC of the telomeric ds-ss junction. The amino acid residues composing the POT-hole show full sequence identity with telomeric proteins from diverse eukaryotes, including *Caenorhabditis elegans* POT-1. The current study builds on this SELEX method, developing an extensive analysis pipeline for SELEX datasets sequenced by next-generation sequencing and achieving a deeper analysis of the resulting sequences. We validated our approach by applying it to the DNA-binding domain of hPOT1, yielding results consistent with a previous SELEX study. Furthermore, we employ our pipeline to characterize the DNA-binding activity of *C. elegans* proteins that are considered homologs of hPOT1: POT-1, POT-2, POT-3, and MRT-1. Our analysis suggests that all four proteins show a binding preference for G-enriched DNA sequences, with POT-1 additionally binding secondary structural elements. Overall, we present a bioinformatics pipeline that is accessible and applicable for determining the nucleic acid-binding properties of a variety of proteins.

## INTRODUCTION

The need to distinguish chromosome ends from DNA damage is the essence of the end-protection problem^1,2^. In mammals, this evasion occurs via the shelterin protein complex, which binds to telomeric DNA at chromosome ends, where there is a double-stranded (ds) region consisting of the repeating sequence TTAGGG-3’/AATCCC-5’ and a single-stranded (ss) G-rich overhang^2^ (**Figure 1a,b**). The kinases ATM and ATR respond to DNA damage. In particular, the 9-1-1 complex-containing DNA clamp/clamp-loader complex recognizes ds-ss junctions, which can form at double-strand breaks (DSBs), and stimulates ATR^3^. The presence of ssDNA is a typical feature of DNA structures that activate ATR kinase signaling, with both RPA-associated ssDNA and a ds-ss junction being important for ATR activation. As mammalian telomeres have naturally occurring ds-ss junctions and ssDNA overhangs, determining how such structures avoid recognition as sites of DNA damage is central to the understanding of genome stability^1,2^.

**Figure 1.**
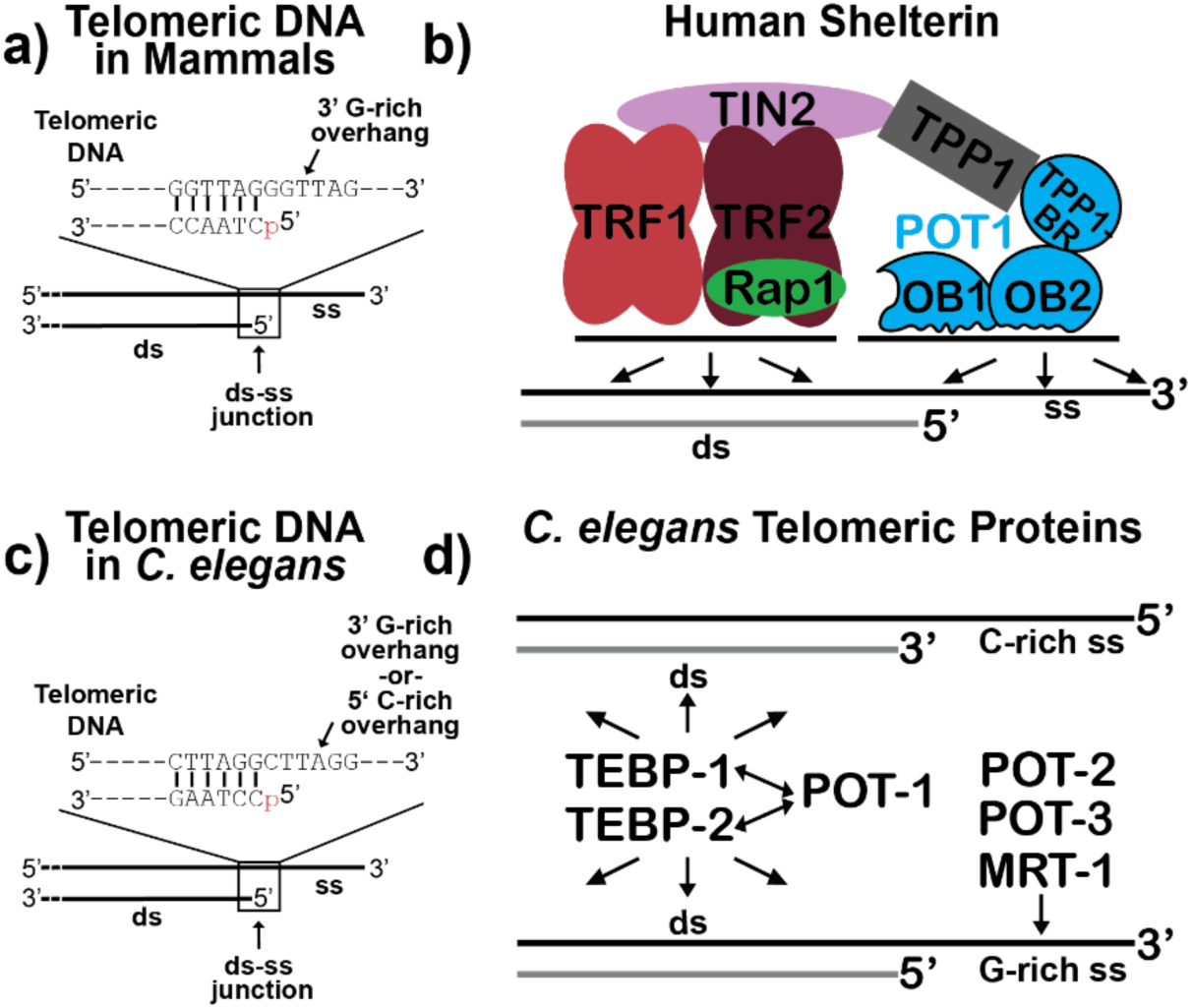
Models of the Structure of Mammalian and *C. elegans* Telomeres^2,61,4,16,15,62,17,22,23,21,20,24^. (a) Mammalian telomeric DNA^2,4,61^. Note the presence of a 5’ phosphate (“p” in red) at the ds-ss junction. (b) The six-member human shelterin complex^2^. (c) Same as (a) but showing a model of *C. elegans* telomeric DNA^15–17,62^. (d) Telomeric proteins of *C. elegans*^16,17,20–24^. Abbreviation: TPP1-BR, TPP1-binding region of hPOT1 protein.

Mammalian POT1, which binds the telomeric ssDNA overhang, inhibits ATR activation at telomeres, and different models have been proposed for how this occurs^2^. According to one proposed mechanism, binding of POT1 to the overhang excludes RPA from binding and thereby inhibits ATR signaling. Yet it was unknown whether any telomeric protein shielded the telomeric ds-ss junction from recognition by factors of the DNA damage response^4^.

Reexamining the DNA-binding properties of human POT1 (hPOT1) that Choi *et al.* determined through systematic evolution of ligands by exponential enrichment (SELEX) brought much more clarity to this question (**Figure 2**)^5,4^. SELEX is an *in vitro* technique for capturing nucleic acid sequences that bind to a target of interest^6–9^. In this procedure, random sequences within a population are interrogated for their binding ability, typically involving the enrichment of these species through a recursive process of selection. SELEX is a well-established technique that has helped characterize the dsDNA-binding properties of the mammalian telomeric protein TRF1^9^ and the transcription factor NF-κB^10^, as well as the ssDNA-binding activity of the phage protein g5p^11^. In another application, SELEX allows for the discovery of nucleic acid sequences (so-called aptamers) that can function as inhibitors of a protein target^7,8^. In their SELEX experiment, Choi *et al.* exposed POT1 to a library of random ssDNA sequences, and after multiple rounds of selection, protein-bound sequences were sequenced and analyzed for motifs^5^. While one category of enriched sequences (“Class I”) seemed to correspond to a known binding site of POT1 (5’-TTAGGGTTAG-3’)^12^, a larger group of sequences (“Class II”) appeared to contain a non-telomeric sequence adjacent to telomeric motifs (**Figure 3a,b**)^5^. Subsequent examination of these results by Tesmer *et al.* suggested that the apparently non-telomeric sequence formed a hairpin that positioned a 5’-phosphorylated C residue adjacent to the region of ss telomeric sequence^4^. A crystal structure of the DNA-binding domain (DBD) of hPOT1 bound to a DNA oligonucleotide resembling the SELEX hit revealed an electropositive concave surface (named the “POT-hole”) comprising four amino acid residues that accommodate the 5’ phosphorylated dC of the ds-ss telomeric junction (**Figure 3c,d**)^4^. This observation agrees with previous studies identifying ATC-5’ as the predominant ds-ss telomeric junction in human cells^13^ and suggests that the POT-hole either promotes formation of this end or, more conservatively, protects it once formed^4^. Mutating POT-hole residues diminished ds-ss junction binding and impaired suppression of DNA damage signaling at telomeres, even when preserving telomeric ssDNA-binding. Importantly, Tesmer *et al.* noted that the POT-hole residues were fully conserved across POT1 homologs in the ciliate *Sterkiella nova* (TEBPα) and *Caenorhabditis elegans* (POT-1), organisms with ds-ss telomeric junctions ending in a phosphorylated dC (*S. nova*^14^; *C. elegans*^15^), suggesting broad conservation of this structural feature and telomeric ds-ss junction binding function among eukaryotes^4^.

**Figure 2.**
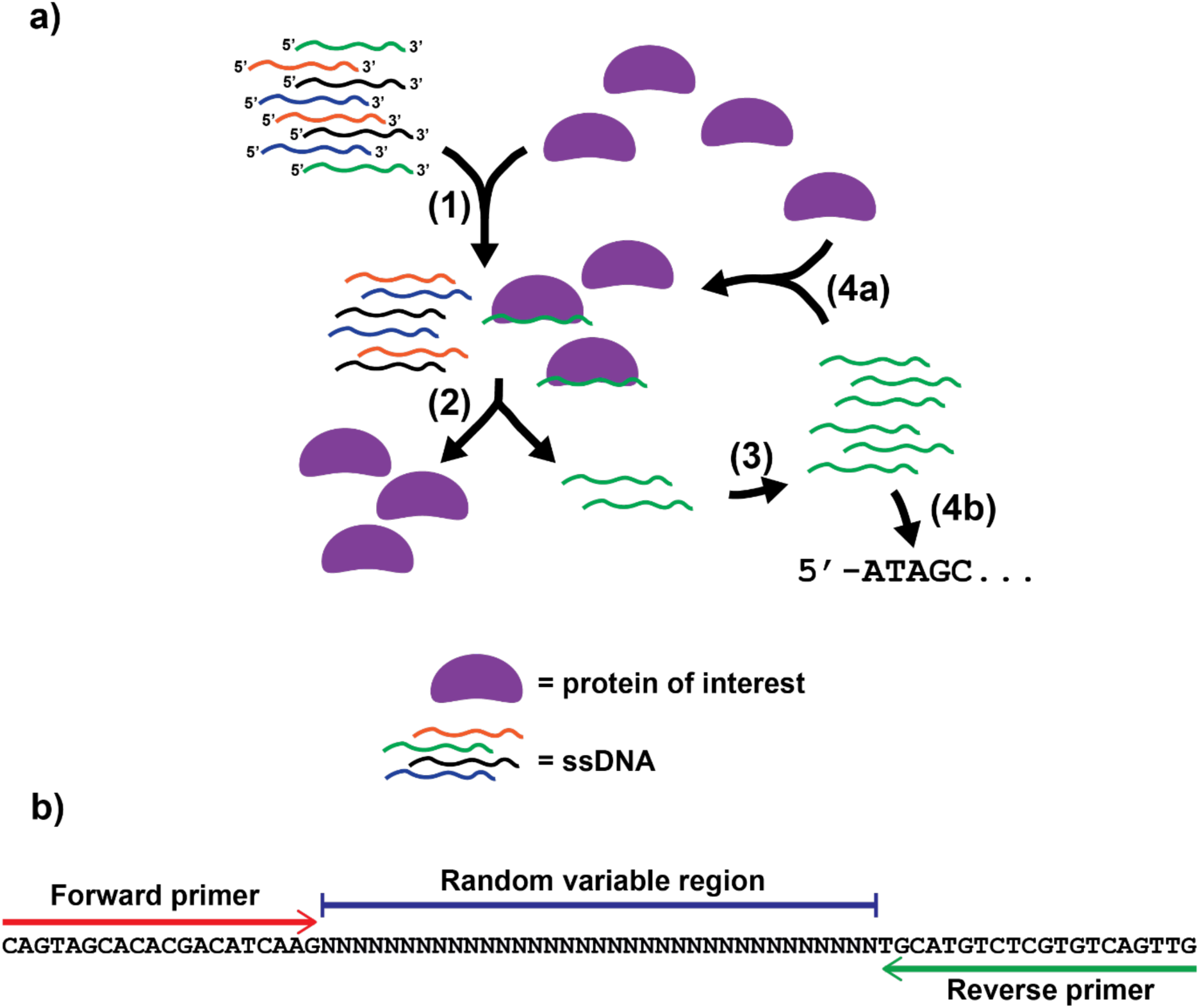
Overview of SELEX with ssDNA as performed by Choi *et al*^5^. (a) In step 1, the protein of interest and unselected ssDNA library are combined and allowed to interact. In step 2, bound ssDNA is freed, and in step 3, the DNA is amplified by PCR, followed by additional selection (4a) or sequencing (4b). (Asymmetric PCR was used in step 3 to form ssDNA for additional cycles of selection.) (b) Schematic of ssDNA oligos used for SELEX.

**Figure 3.**
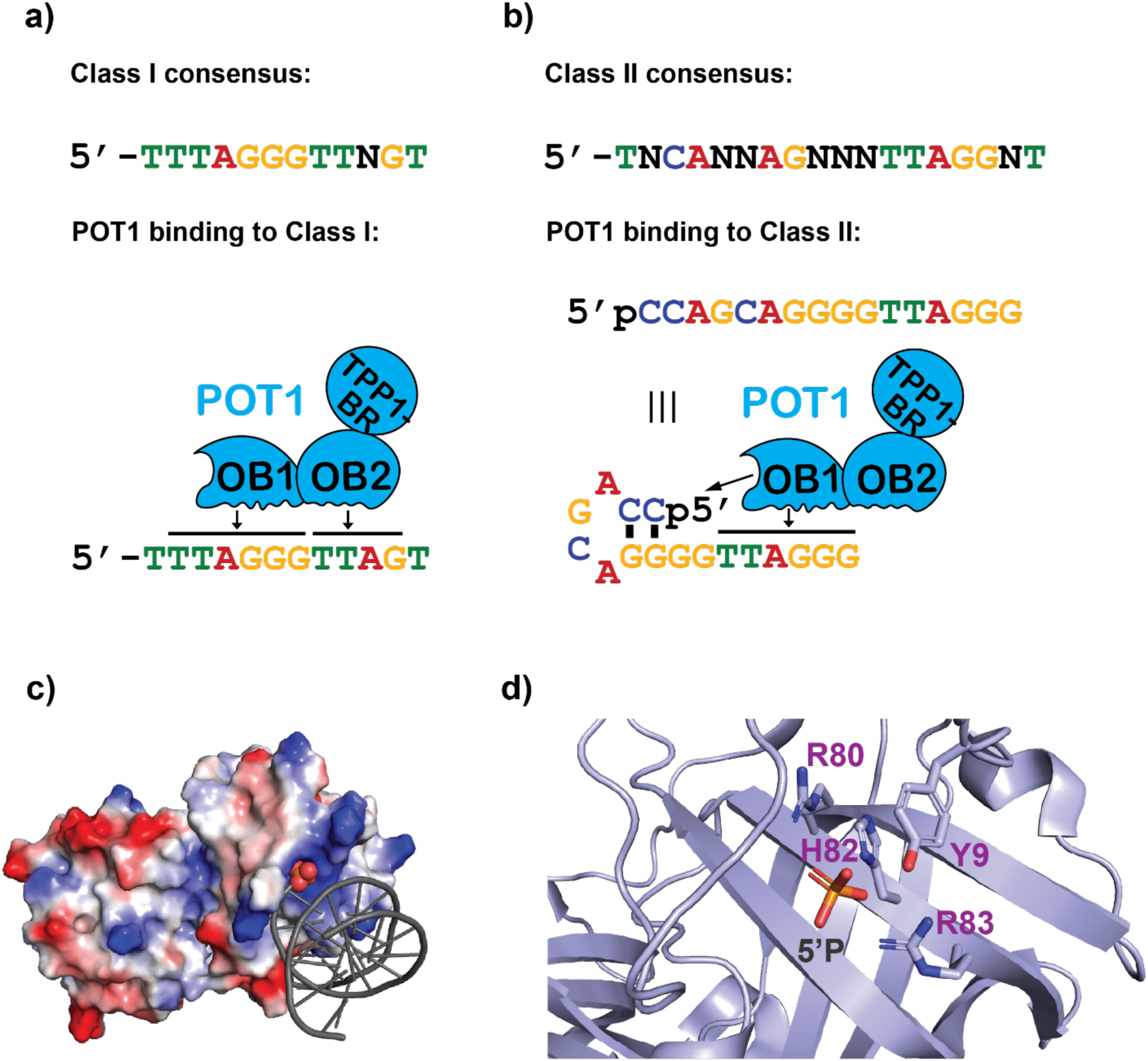
Human POT1 Binds to Telomeric ssDNA and to a ds-ss Junction^12,5,4^. (a) Class I consensus sequence from the SELEX experiment by Choi *et al.* (top) and the proposed manner of DNA-binding by POT1 (bottom)^5^. (b) Class II consensus sequence from the SELEX experiment by Choi *et al.* (top) and a model of DNA-binding by POT1 based on these results and those of Tesmer *et al*^5,4^. (c) Electrostatic surface representation of hPOT1 DBD bound to the telomeric ds-ss DNA junction showing charge complementarity of the POT-hole and the 5’ end of the chromosome (PDB: 8SH1)^4^. Red and blue indicate electronegative and electropositive regions, respectively. DNA is shown in gray with atoms of the 5’ phosphate group depicted as spheres. (d) Close-up of the POT-hole with the side chains of its residues displayed in a stick representation. The 5’ phosphate is shown, but the rest of the DNA substrate is hidden for clarity. Diagrams in (c) and (d) were generated using PyMOL Molecular Graphics System, Version 3.1.6.1 Schrödinger, LLC.

Like mammalian telomeres, the telomeres of *C. elegans* are thought to contain a region of dsDNA followed by a ssDNA overhang (**Figure 1c**)^2,15,16^, but the telomeric repeat sequence differs at one position within the hexad (human: TTAGG<u>G</u>^2^, *C. elegans*: TTAGG<u>C</u>^15,16^). Unlike telomeres from all other characterized eukaryotes, including mammals, which are only known to contain a 3’ G-rich ssDNA overhang, one study suggests that, in addition to ending in a 3’ G-rich overhang, *C. elegans* chromosomes may also end in an overhang on the 5’ C-rich strand^1,17,16^.

As in mammals^2^, *C. elegans* responds to DNA damage through the kinases ATM and ATR^18,19^, the latter of which is activated by 9-1-1^3^. Disruption of ATR in *C. elegans* impairs cell cycle arrest and apoptosis in response to ionizing radiation (IR)^18^, which generates double-strand breaks (DSBs)^19^. The telomeric DNA of *C. elegans* is bound by an assembly of proteins that bears structural and functional resemblance to shelterin (**Figure 1d**)^16,20,21,2^. In particular, *C. elegans* telomeres contain two telomeric dsDNA-binding proteins, TEBP-1 and TEBP-2^22,23,16^, echoing the manner in which TRF1 and TRF2 bind to dsDNA in shelterin^2^. Four telomeric proteins in *C. elegans* are considered homologs of hPOT1: POT-1, POT-2, POT-3, and MRT-1^16,20^. Interestingly, these telomeric proteins are reported to have varying ssDNA-binding preferences, with POT-1 exhibiting sequence selectivity for the putative C-rich 5’ overhang and POT-2, POT-3, and MRT-1 favoring the G-rich 3’ overhang^24,17,20^. Furthermore, while mammalian POT1 proteins bind to TRF1 and TRF2 indirectly via the shelterin subcomplex TIN2-TPP1^2^, *C. elegans* POT-1 binds to TEBP-1 and TEBP-2 directly^16,22,23^. This direct bridging of different DNA-binding proteins at chromosome ends qualifies *C. elegans* as a suitable model for dissecting shelterin function.

Expanding on the approach that Choi *et al.* applied to characterize hPOT1 (in which they cloned into plasmids and sequenced fifty sequences after the final round of SELEX)^5^, we employed an *in vitro* protocol that combined SELEX with high-throughput, next-generation sequencing (NGS)^25^ and developed a bioinformatics pipeline to identify enriched DNA-protein interaction motifs. Using this method, we determined the DNA-binding preference of hPOT1 as a proof of principle and investigated the DNA-binding properties of the *C. elegans* proteins POT-1, POT-2, POT-3, and MRT-1. Based on these efforts, we present a method for HT-SELEX analysis that does not require significant coding experience and that is accessible on a Windows operating system (OS), making it broadly usable by the biomedical research community.

## RESULTS

Our approach to determine the DNA-binding properties of *C. elegans* telomeric proteins expanded on the SELEX technique that Choi *et al.* had used to study the hPOT1^5^ by subjecting the selected DNAs to high-throughput sequencing, a method known as HT-SELEX^25,26^. We reasoned this to be a useful, unbiased assay for characterizing protein-DNA interactions. Such an approach has several advantages compared to standard SELEX. (1) Being compatible with Illumina adaptor-based sequencing, it is high-throughput, allowing for the sequencing of a larger portion of the selected DNAs compared to Sanger sequencing. (2) It does not involve molecular cloning. (3) Finally, it is economical (∼$60 per sample for ∼50,000 reads) through services such as Amplicon-EZ NGS (Azenta).

Towards our goal to develop HT-SELEX for telomeric ssDNA-binding proteins, we created an analysis pipeline for processing the high-throughput NGS data from the DNAs selected in the SELEX procedure and finding motifs that were enriched for binding a protein of interest. An initial step was to survey existing computational tools available for SELEX data analysis and consider their appropriateness for our application. Previously developed analysis tools included APTANI2^27,28^, FSBC^29^, FASTAptameR 2.0^30,31^, Galaxy Project^32,33^, DeepSELEX^34^, Inimotif^35^, AptaSUITE^25,36^, and the MEME Suite^37^.

APTANI2 has the advantage of scoring SELEX-enriched sequences based on the abundance of certain motifs and the stability of secondary structures predicted to form in the molecules^28^. Moreover, APTANI2 can be accessed via a graphical user interface (GUI) for ease of use. However, while the method for predicting secondary structures appears to be intended for RNA^28,38^, it is unclear whether it would be valid for SELEX performed with DNA. On the other hand, Kato *et al.* present FSBC as a tool for grouping SELEX-enriched nucleic acids based on commonly appearing sequences, allowing researchers to validate these candidate, target-binding groups experimentally^29^. However, detailed instructions or a manual for the FSBC analysis tool were not readily available.

FASTAptameR 2.0 can be used to find motifs in either RNA or DNA, employing FSBC in this process^31^. Yet, while Kramer *et al.* present FASTAptameR 2.0 as a useful tool for an initial search for motifs, they highly recommended the MEME Suite^37^ (discussed in more detail below) and other resources for motif identification^31^. Thus, FASTAptomerR 2.0 may not be the optimal tool to interpret our SELEX data for determining the DNA-binding preference of a protein^31^. The Galaxy Project offers a set of straightforward online tools that can be employed for HT-SELEX data analysis^32^.

However, Thiel and Giangrande acknowledge that the Galaxy Project may not be well suited for applications requiring more thorough investigation of motifs. (In fact, one of the sources that they cite after this statement is regarding FASTAptamer^30^, an earlier version of FASTAptameR 2.0^31,32^.) This suggests that the Galaxy Project tools may not be optimal for our purposes^32^. An alternate option, DeepSELEX is notable for its application of neural networks to interpret HT-SELEX results^34^; however, we sought a tool that could function on a Windows OS, while DeepSELEX functions best in a Linux context^39^. The Inimotif tool shows promise in light of its use of Hamming distance (i.e., the number of nucleotide changes needed to change an observed sequence into another observed sequence used as a reference, either in the forward or reverse orientation) to judge the similarity between sequences enriched in the SELEX experiment^35^. This enables researchers to quantify and visualize the relative abundance and sequence-similarity of highly enriched sequences. However, detailed information for using this analysis tool is not available to the best of our knowledge. Also, Inimotif requires Python, and potentially, a non-trivial amount of coding expertise^40^.

AptaSUITE^36^ is a computational toolkit for analysis of HT-SELEX data that includes AptaPLEX^25^, AptaCLUSTER^41^, and AptaTRACE^42^. Users of AptaPLEX can merge paired-end reads and identify and isolate the randomized region of the sequenced DNAs from SELEX^25^. Additionally, AptaCLUSTER allows researchers to group SELEX-enriched sequences into categories based on sequence similarity. Such a tool appeared suitable to find motifs by grouping similar sequences. Moreover, AptaTRACE enables researchers to query HT-SELEX results for motifs based on nucleotide sequence and secondary structure^42^. An important insight from Dao *et al.* is that sequences that preferentially bind will also be in the setting of a secondary structure that favors binding. Over the course of selection, one would expect the enriched sequences to increasingly coincide with secondary structures that facilitate interaction with the target. By sequencing and analyzing the products of different rounds of the SELEX procedure, researchers can track the selection of nucleotide sequence motifs and their structural settings. Although the algorithm is intended to be used for both RNA and ssDNA, it is not clear that the secondary structural predictions that are at the core of AptaTRACE are optimized for ssDNA to the extent that they are for RNA. A major advantage of the AptaSUITE is its user-friendly GUI, allowing users to process, analyze, and visualize their SELEX data on a Windows-based system^41,42,25,36^.

Finally, the MEME Suite offers a variety of analysis tools for DNA, RNA, and protein motifs^37^, including *de novo* motif identification^43–47^, detection of known motifs in a set of sequences^45,48–50^, and comparison of motifs to one another^44,51^. Two of these tools that were of particular interest for finding motifs *de novo* were the suite’s namesake tool, MEME^43^, and another algorithm capable of handling larger datasets known as STREME^47,37^. There were several advantages to using STREME. To begin, users of the algorithm could easily submit the HT-SELEX results as a FASTA file for analysis. Secondly, in one submission of SELEX results to STREME, we could query our dataset for motifs of various sizes (e.g. 10-25 nt motifs). Thirdly, the STREME output showed the abundance of each discovered motif in the dataset and could provide an estimate of statistical significance for each motif that accounted for multiple comparisons when several motifs were identified. In addition, detected motifs were displayed as sequence logos that present the information content at each nucleotide position by the height of each stack of letters, while the relative heights of letters in a given stack indicate the relative abundance of each nucleotide at that site^37,47^. In sequence logos, information content is quantified in bits and measures how important the different nucleotide positions of a motif are^52^. Given that there are four possible nucleotides at any position, two bits are the maximum information content at any position, as would be seen if exactly one nucleotide (e.g., always T and never A, C, or G) were observed at that site. Unlike a consensus sequence, which only shows the predominant residue at each position, logos depict heterogeneity of nucleotides at each site. Furthermore, the letters at each position are stacked from top to bottom in decreasing order of abundance at that site.

The advantages and disadvantages of various analysis tools are summarized in **Table 1**. In light of the available analysis tools, our strategy was to integrate AptaSUITE^36^ and STREME^47^ into a pipeline for *de novo* motif identification in our HT-SELEX data. After SELEX, DNAs undergo high-throughput sequencing, yielding paired-end reads that are imported into AptaSUITE^36^ as FASTQ files. The AptaPLEX algorithm merges paired-end reads and extracts variable randomized regions from the pool of sequenced DNAs^25^. We exported this processed data from AptaSUITE as a FASTA file^36^ and used a text editor (Notepad++) to add a unique identifier for each sequence (which was a requirement prior to submitting the dataset to STREME^37,47^). STREME analysis was performed using the MEME Suite public web server^37,47^. Although the oligonucleotides in the SELEX experiment were ssDNA in the selection process, the STREME algorithm presents the motifs it identifies as dsDNA motifs, giving the sequence logo of one strand as well as the reverse complement. A function in R was used to determine the orientation of the discovered motifs in the dataset by computing and graphing the incidence of a motif and its reverse complement.

**Table 1.** Summary of Analysis Tools for SELEX Data.

| Analysis Tool | Advantages | Disadvantages |
| --- | --- | --- |
| **APTANI2**<sup>27,28,38</sup> | Evaluates sequences in terms of abundance of motifs and the secondary structures associated with them<br>Has a GUI | Unclear if secondary structural predictions are suitable for DNA |
| **FSBC**<sup>29</sup> | Groups sequences based on recurring motifs | No manual available, to the best of our knowledge |
| **FASTAptamerR 2.0**<sup>30,31</sup> | Adaptable to DNA and RNA | May not be appropriate for definitive motif identification |
| **Galaxy Project**<sup>32</sup> | Ease of use | May not be appropriate for definitive motif identification |
| **DeepSELEX**<sup>34,39</sup> | Employs neural networks | More appropriate for Linux vs. Windows |
| **Inimotif**<sup>35,40</sup> | Applies Hamming distance for comparison of enriched sequences | No detailed manual available, to the best of our knowledge<br>May require familiarity with Python |
| **AptaSUITE**<sup>25,36,37,41,42,47</sup> | AptaPLEX facilitates processing of NGS data<br>AptaCLUSTER groups sequences based on similarity<br>AptaTRACE identifies motifs while accounting for sequence and 2° structure<br>User-friendly GUI<br>Suitable for Windows OS | AptaTRACE may not be well suited for predicting DNA secondary structure<br>Sequences in FASTA files exported from AptaSUITE require unique identifiers before submission to STREME |
| **STREME**<sup>47</sup> (part of the MEME Suite<sup>37</sup>) | Datasets can easily be submitted as FASTA files<br>Able to seek motifs of varying length in one search<br>Detailed statistical analysis of motifs identified<br>Sequence logos of motifs | STREME treats a dataset submitted as ssDNA as if it were dsDNA |

Our SELEX procedure began with an ssDNA library (hereafter referred to as the “N35 library”) with the same constant regions corresponding to forward and reverse primers and a central 35-nucleotide variable region as employed for hPOT1 by Choi *et al.* previously (**Figure 2b**)^5^. To see the baseline composition of the variable region prior to SELEX, we subjected the unselected N35 library to NGS. If all four nucleotides (A, T, G, C) had equal probability at each position in the variable region, then one would expect a frequency of 25% for each nucleotide throughout the region. Based on the sequencing results, most positions showed nucleotide frequencies in the 20-33% range, with a slight overall preference for A and T compared to G and C (**Figure 4**)^36,25^. Thus, the N35 library did not appear to have significant bias in nucleotide composition prior to selection.

**Figure 4.**
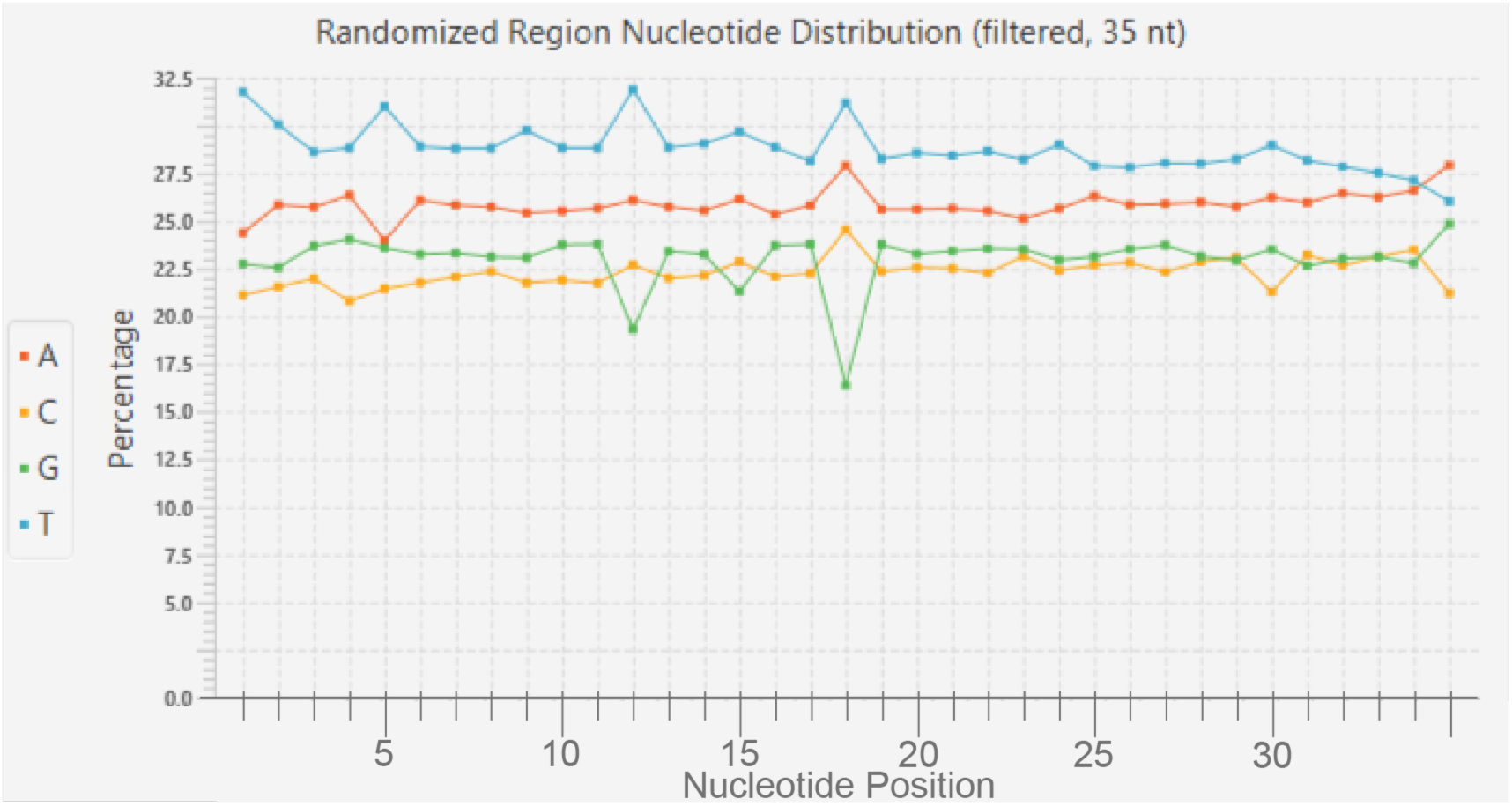
Composition of the unselected N35 library, adapted from AptaSUITE^36,25^.

Furthermore, as proof of principle for the ability of our HT-SELEX procedure to reveal the binding properties of a protein of interest, we performed SELEX with the N35 library and purified hPOT1 DNA-binding domain containing a polyhistidine-SUMOstar tag (hereafter called hPOT1-DBD). If hPOT1-DBD were to bind to a specific motif in the ssDNA population, one would expect the composition of the variable regions in the SELEX-enriched sequences to change significantly from that of the unselected library. Indeed, analysis of DNA sequenced from the fourth round of SELEX showed a markedly different landscape in the variable region (**Figure 5**)^36,25^. Firstly, instead of being roughly parallel to each other throughout the 35 nt window, the frequencies of the four nucleotides fluctuate with clear peaks and troughs. Secondly, the top two most frequent nucleotides in most nucleotide positions are G and T, fully consistent with hPOT1-DBD binding the TTAGGG human telomeric repeat with high sequence specificity^2^. Inspecting regions of the plot more closely, it can be noted that positions 24-33 are relatively depleted of C residues and enriched for G. This observation suggests that the nucleotide composition of the variable region has evolved due to the binding of particular sequences to hPOT1-DBD and their resulting enrichment in the population.

**Figure 5.**
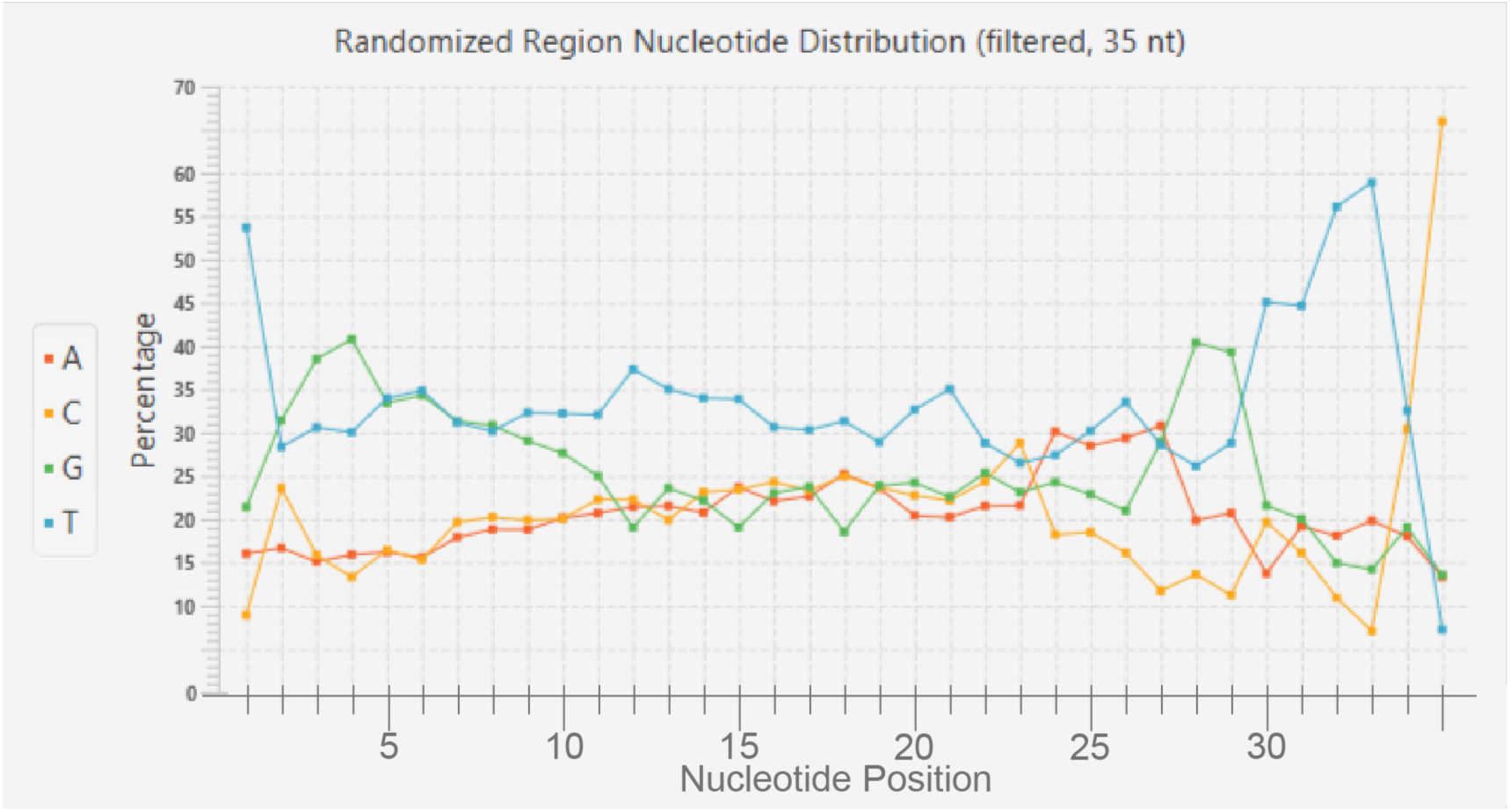
Nucleotide composition of the variable region after the fourth round of SELEX using hPOT1-DBD, adapted from AptaSUITE^36,25^.

In addition, we subjected the hPOT1-DBD HT-SELEX dataset to motif identification via the STREME algorithm (**Figure 6**)^37,47^. The results showed similarity to the DNA binding sequence specificity reported by Choi *et al.* with FLAG-tagged hPOT1 protein^5^, with the top six motifs (in terms of abundance, i.e., the percentage of oligonucleotide sequences containing them) including three (motifs 1.4-1.6) with consensus sequences that contained all or most of the mammalian telomeric repeat TTAGGG (**Figure 6a**)^2^. In fact, the consensus of Motif 1.6 contains the first eight bases (5’-TTAGGGTT-3’) of the ten-nucleotide POT1-binding site that Choi *et al.* identified in their Class I SELEX result^5^. Importantly, Motif 1.1 contains an apparent hairpin-forming region^53,54^ that resembles the Class II SELEX hit that Choi *et al.* discovered^5^, and which later investigation would strongly suggest could form a hairpin simulating a telomeric ds-ss junction (**Figure 6b**)^4^. In the SELEX experiment by Choi *et al.*, Class II was the predominant species identified^5^, and likewise, in our experiment, Motif 1.1 had the highest abundance (present in 28.4% of sequences). Thus, comparison of our hPOT1-DBD results with SELEX results of previous work validate our use of this procedure to characterize DNA-binding activity of proteins. This paves the way for us to employ the same method to elucidate DNA-binding properties of *C. elegans* proteins that are considered homologs of hPOT1: POT-1, POT-2, POT-3, and MRT-1^16^.

**Figure 6.**
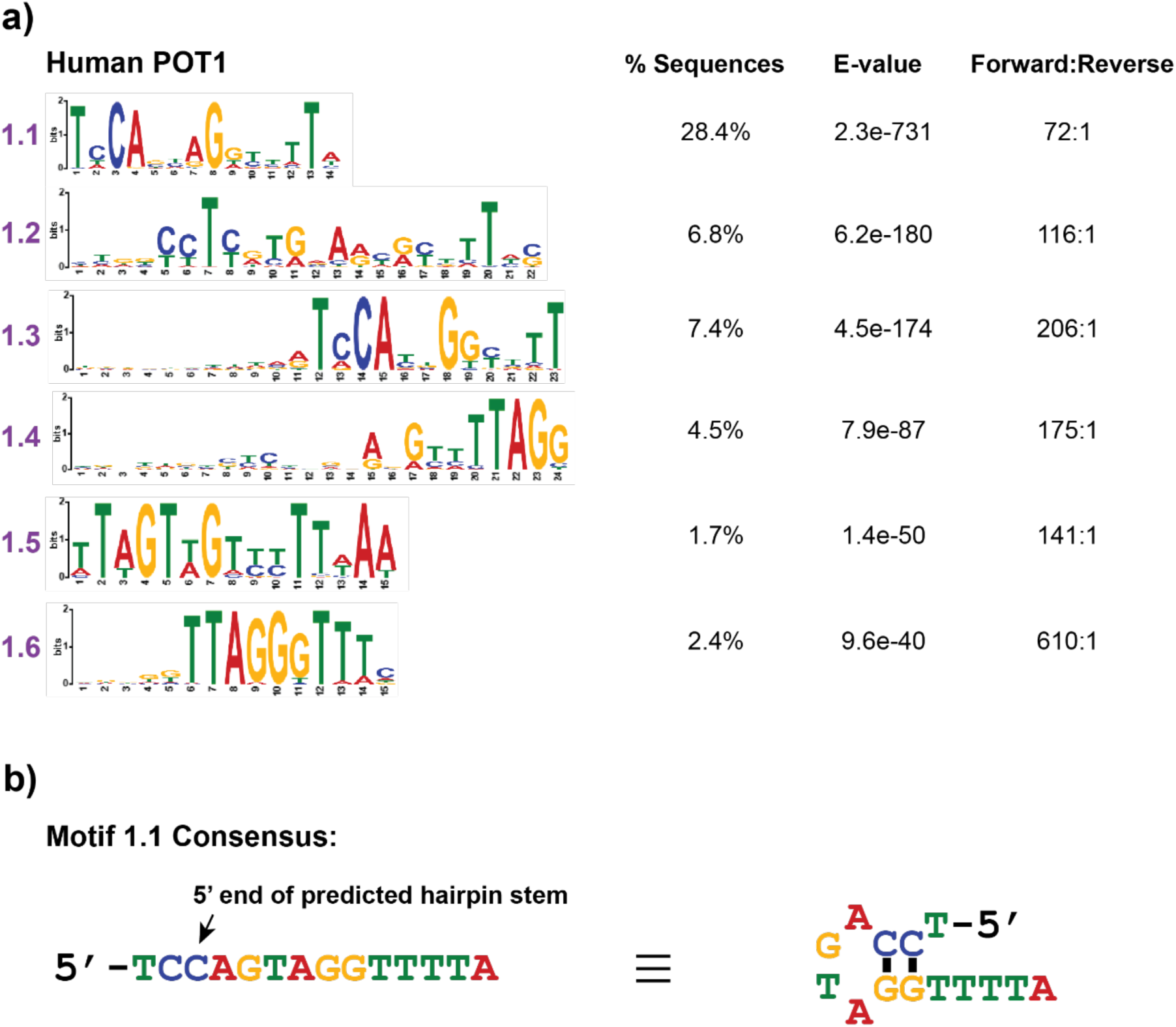
HT-SELEX with hPOT1-DBD. (a) Analysis of results showing the six most abundant motifs (listed in decreasing E-value). % Sequences indicates the proportion of sequences containing the motif, E-value is an adjusted measure of statistical significance, and Forward:Reverse indicates the relative abundance of the motif in either orientation. See **Methods** section for details regarding these metrics. (b) Consensus sequence of Motif 1.1 and the secondary structural prediction (using the UNAFold web server^53,57,58^).

While the DNA-binding region of hPOT1 contains two oligonucleotide/oligosaccharide (OB) folds (OB1 and OB2), which bind to telomeric ssDNA^12^, only one OB fold has been identified in each of the four *C. elegans* proteins (**Figures 7,8**)^16,17,24^. The OB fold of POT-1 resembles OB1, while the OB folds of POT-2, POT-3, and MRT-1 resemble OB2^16,17,24^.

**Figure 7.**
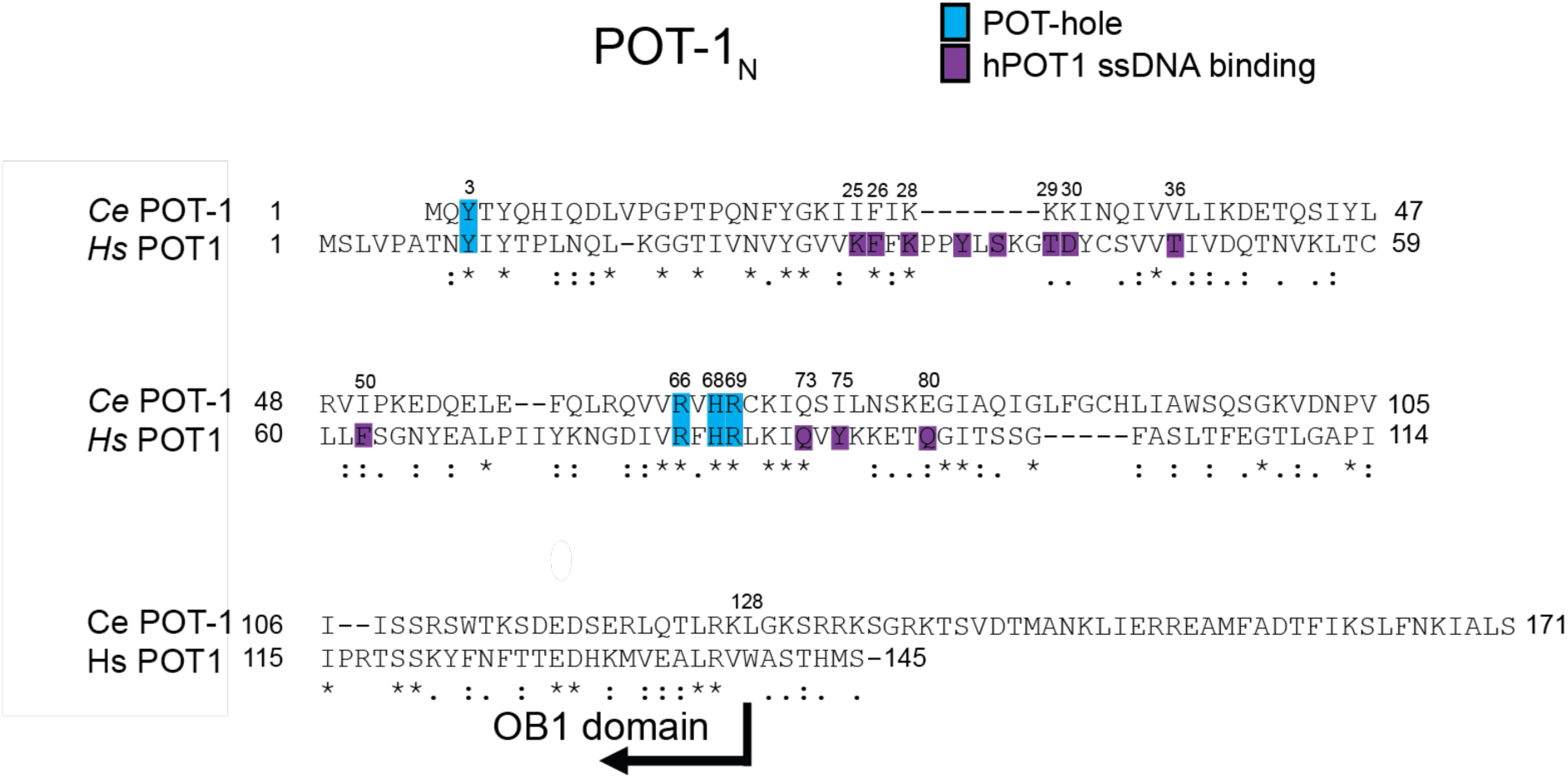
POT-Hole Residues are Identical Between Human POT1 and *C. elegans* POT-1. Amino acid sequence alignment of OB1 of hPOT1 and *Ce*POT-1 proteins, made using Clustal Omega^63^. An asterisk indicates identical residues, two dots indicate substantially similar residues, and one dot indicates modestly similar residues at a given position.

**Figure 8.**
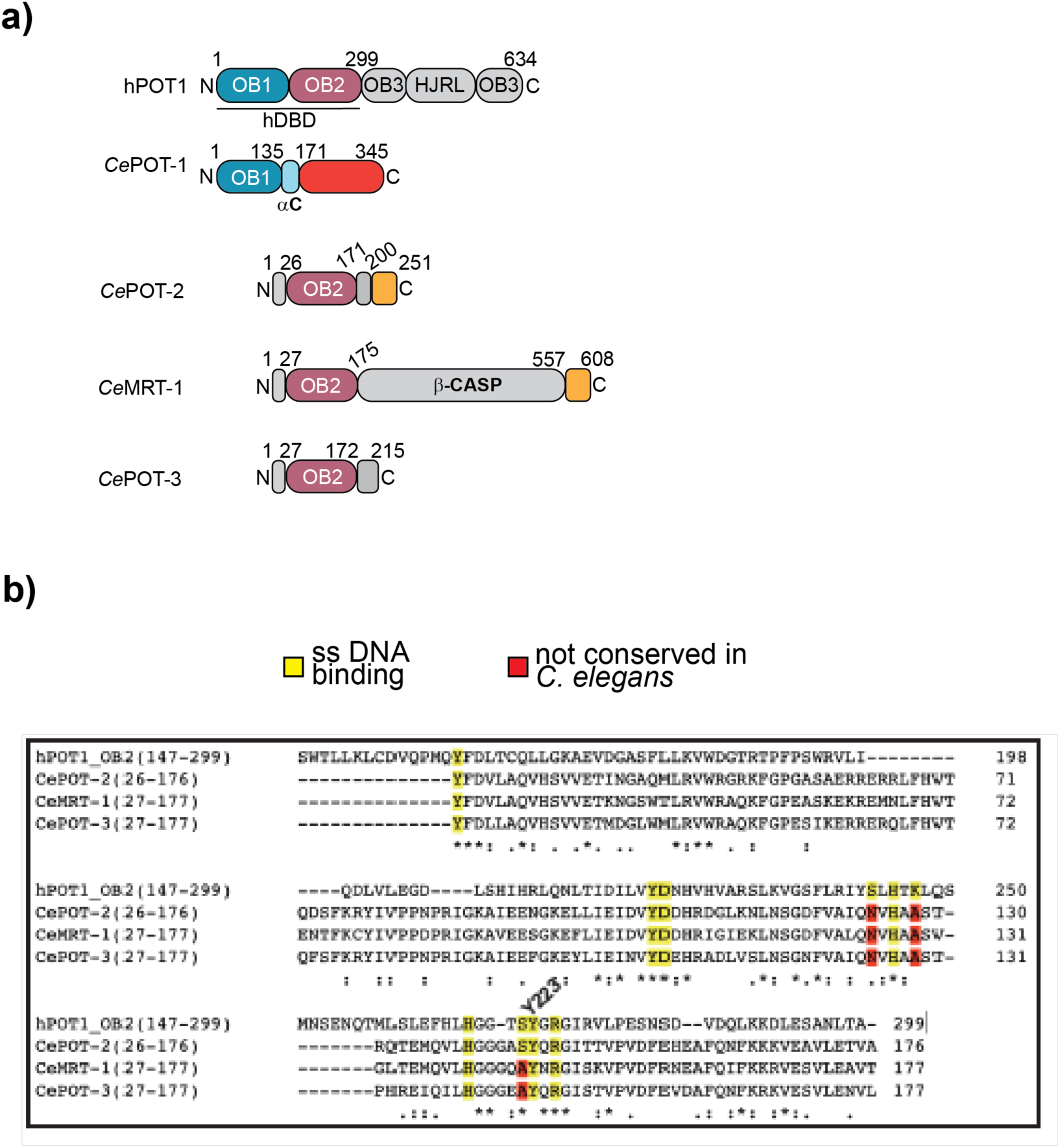
Human and *C. elegans* Telomeric ssDNA-Binding Proteins^2,17,20,24^. (a) Schematic of human and *C. elegans* (*Ce*) proteins with amino acid numbering. (b) Amino acid sequence alignments of hPOT1 OB2 with the *C. elegans* proteins POT-2, MRT-1, and POT-3, generated using Clustal Omega^63^. Below each alignment, an asterisk indicates identical residues, two dots indicate substantially similar residues, and one dot indicates modestly similar residues at a given position. Note that the full C-terminal region of hPOT1 is not included in the alignment. Abbreviations: N, N-terminus; C, C-terminus; HJRL, Holliday junction resolvase-like^2^; αC, α-helix C; β-CASP, metallo-β-lactamase-associated CPSF-Artemis-SNM1/PSO2^24^.

Intriguingly, *C. elegans* POT-1 shares the same four POT-hole residues present in hPOT1, raising the question of whether it likewise binds the 5’-phosphorylated dC of the telomeric ds-ss junction (**Figure 7**)^4^. However, strikingly, it lacks equivalence for residues that form the ssDNA-interface with hPOT1. In a recently submitted manuscript, we demonstrated that the ssDNA-binding surface of POT-1 instead binds its partners at telomeres, TEBP-1 and TEBP-2^16,22,23^, resulting in a tight protein complex that is essential for POT-1 protein stability (Tesmer *et al., submitted*). In the study described in the current manuscript, we performed HT-SELEX using a SUMO-tagged, N-terminal region of POT-1, which contains the apparent POT-hole residues^4^ (POT-1_N_), co-purified in complex with the polyhistidine-SUMO-tagged POT-1 binding domain (PBR) of TEBP-1, hereafter called “POT-1_N_-TEBP-1_PBR_” (**Figure 9**)^22^. The inclusion of the TEBP-1 fragment was essential to stabilize the POT-1 fragment (Tesmer *et al., submitted*). In contrast with the observations with hPOT1-DBD, none of the top six motifs identified by STREME had a consensus sequence containing a full *C. elegans* telomeric repeat (TTAGGC)^15,16^. Furthermore, none of the other fourteen discovered motifs had a full repeat in their consensus (data not shown). On the other hand, three of the six most abundant motifs (motifs 2.1, 2.4, and 2.5) were enriched for a string of three or more contiguous G nucleotides, a nucleotide that was somewhat underrepresented in the unselected N35 library (**Figure 4**), with many of these positions in the sequence logos showing high information content (**Figure 9**). Two other motifs (motifs 2.2 and 2.6) also showed G-enrichment, although not in a strongly contiguous pattern (**Figure 9**). This G-rich pattern is especially notable given the previous report that POT-1 binds preferentially to the C-rich strand ^17^.

**Figure 9.**
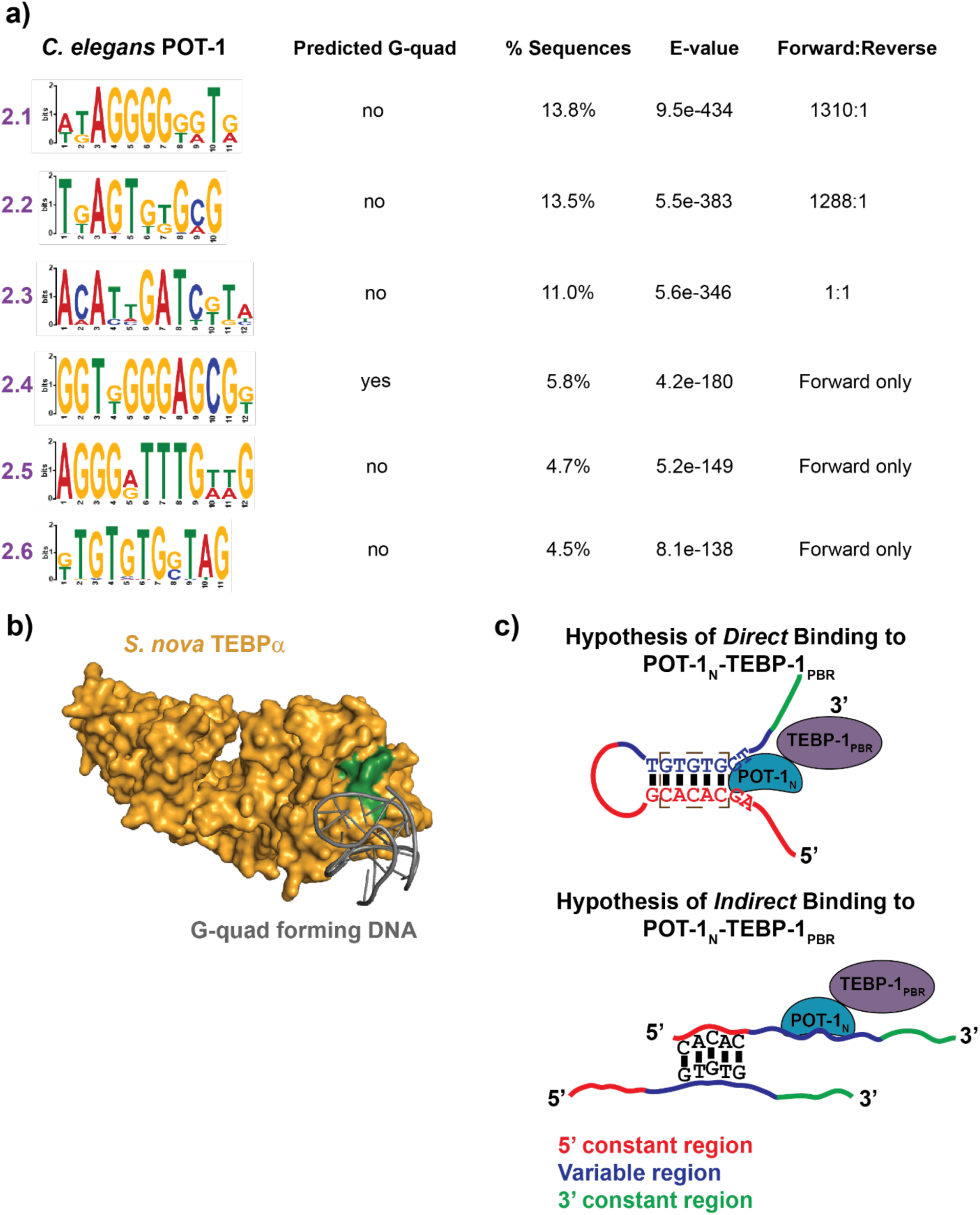
Analysis of HT-SELEX with *C. elegans* POT-1_N_-TEBP-1_PBR_. (a) The six most abundant motifs (listed in decreasing E-value). % Sequences indicate the proportion of sequences containing the motif, E-value is an adjusted measure of statistical significance, and Forward:Reverse indicates the relative abundance of the motif in either orientation. See **Methods** section for details regarding these metrics. (b) Surface model of ssDNA-bound *S. nova* TEBPα (yellow) bound simultaneously to another DNA forming a G-quadruplex, shown in gray (PDB: 1JB7)^56,4^. The ssDNA is not shown, for clarity purposes. Residues corresponding to the POT-hole residues of hPOT1 are shown in green. Image was generated using PyMOL Molecular Graphics System, Version 3.1.6.1 Schrödinger, LLC. (c) Proposed mechanisms for direct and indirect binding to the POT-1_N_-TEBP-1_PBR_ complex of an oligo containing Motif 2.6. Hairpin prediction is based on UNAFold web server^53,57,58^. One of the motifs that had lower abundance (3%) contained 5’-TAGGGT-3’, which is a permutation of the mammalian telomeric repeat (data not shown)^2^.

We found plausible mechanisms for why some of these motifs were enriched. Using a tool for prediction of G-quadruplexes in DNA sequences (QGRS Mapper), we predicted the consensus sequence of Motif 2.4 to form a G-quadruplex, suggesting that *C. elegans* POT-1 may bind to sequences that form such structures^55^. Moreover, G-quadruplex DNA binds to *S. nova* protein TEBPα (considered a homolog of hPOT1) adjacent to four amino acids that are identical to the human POT-hole residues^56,4^, suggesting that a similar mechanism may explain the interaction between *C. elegans* POT-1 and Motif 2.4 (**Figure 9b**). Moreover, Motif 2.6 contains a repetitive 5’-GTGTG-3’ sequence, which is complementary to the 5’-CACAC-3’ sequence of the 5’ constant region of oligos in the N35 library (**Figure 2b**). This suggests a potential mechanism for how Motif 2.6 was enriched. Supported by secondary structure prediction on the UNAFold web server, this model suggests that ssDNAs containing Motif 2.6 form hairpins through intramolecular base pairing between the complementary five-nucleotide sequences (**Figure 9a,c**)^53,57,58^. Perhaps POT-1 binds to the ds-ss junction at the base of the loop in a manner reminiscent of capping of the ds-ss telomeric junction by hPOT1 ^4^. Alternatively, the 5’-GTGTG-3’/5’-CACAC-3’base pairing of Motif 2.6 may be achieved in *trans*, allowing for the variable region of some oligos to anneal to the complementary five-nucleotide sequence in the constant region of other oligos that directly bind POT-1_N_-TEBP-1_PBR_ (**Figure 9c**). Such intermolecular base pairing could allow oligos to “piggyback” on ssDNAs that directly bind protein during the HT-SELEX procedure, undergoing selection due to indirect protein binding. We favor the hairpin model over intermolecular base pairing due to thermodynamic considerations.

Furthermore, we performed HT-SELEX with the *C. elegans* POT-2 protein, which is reported to bind ssDNA with preference for the *C. elegans* ss G-rich strand (**Figure 10**)^17,24^. Whether we searched for motifs 10-25 nucleotides (nts) in length (as we had done for hPOT1 and *Ce*POT-1) or 5-10 nts in length, none of the motifs discovered by the STREME algorithm had a consensus sequence containing a full telomeric repeat sequence (TTAGGC or any permutation thereof)^15,16^. (Given that only one OB fold has been identified in POT-2, POT-3, and MRT-1 proteins^16,17,24^, we wanted to examine a shorter motif length, 5-10 nts, which might accommodate binding of a single OB fold to ssDNA^12^.) However, enrichment of G nucleotides at positions of high information content is a common feature of the most abundant motifs in each case, which is consistent with the preference of POT-2 for G-rich strand telomeric ssDNA reported by others^17,24^.

**Figure 10.**
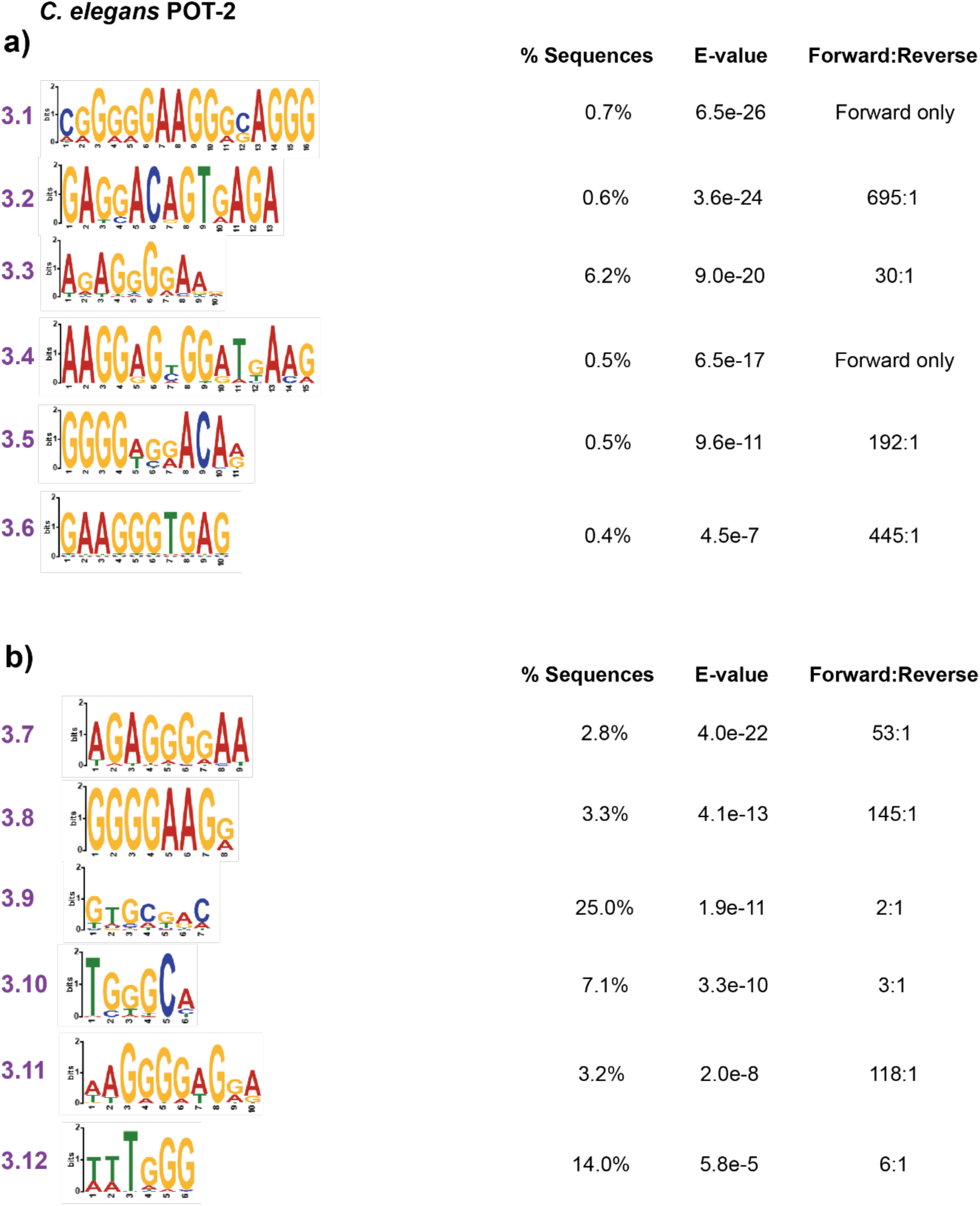
Analysis of HT-SELEX with *C. elegans* POT-2. (a) The six most abundant motifs (listed in decreasing E-value) when searching for motifs 10-25 nts in length. % Sequences indicates the proportion of sequences containing the motif, E-value is an adjusted measure of statistical significance, and Forward:Reverse indicates the relative abundance of the motif in either orientation. See **Methods** section for details regarding these metrics. (b) Same as (a) but showing the results when searching for motifs 5-10 nts in length. After running STREME for POT-2 using either motif range, one of the less abundant motifs when searching for motifs 10-25 nts in length contained 5’-GGTTAGGG-3’ in the consensus sequence and one of the less abundant motifs when searching for motifs 5-10 nts in length contained 5’-TTAGGG-3’, which are examples of the mammalian telomeric repeat sequence (data not shown)^2^. While these were in low abundance (% Sequences ≈ 0.3% or less), perhaps they could be explained by some promiscuity of the POT-2 protein for a different telomeric repeat sequence^15,16^.

Our results for HT-SELEX with *C. elegans* POT-3 tell a similar story to that of POT-2, whether we searched for motifs 10-25 nts or 5-10 nts in length (**Figure 11**). When we searched for 10-25 nt motifs, one of the less abundant motifs (with % Sequences = 0.7%) contained a full permutation of the *C. elegans* telomeric repeat (5’-GGCTTA-3’) in the consensus sequence, although this permutation is different from the 5’-GCTTAG-3’ preference reported previously (data not shown)^20^. For either search, the six most abundant sequences out of the twenty that STREME identified are all G-rich, which is in agreement with the G-rich strand ssDNA binding activity already described for POT-3^20^. The similar results for POT-2 and POT-3 may be explained by the high degree of sequence identity between POT-2 and POT-3 OB domains (**Figure 8b**).

**Figure 11.**
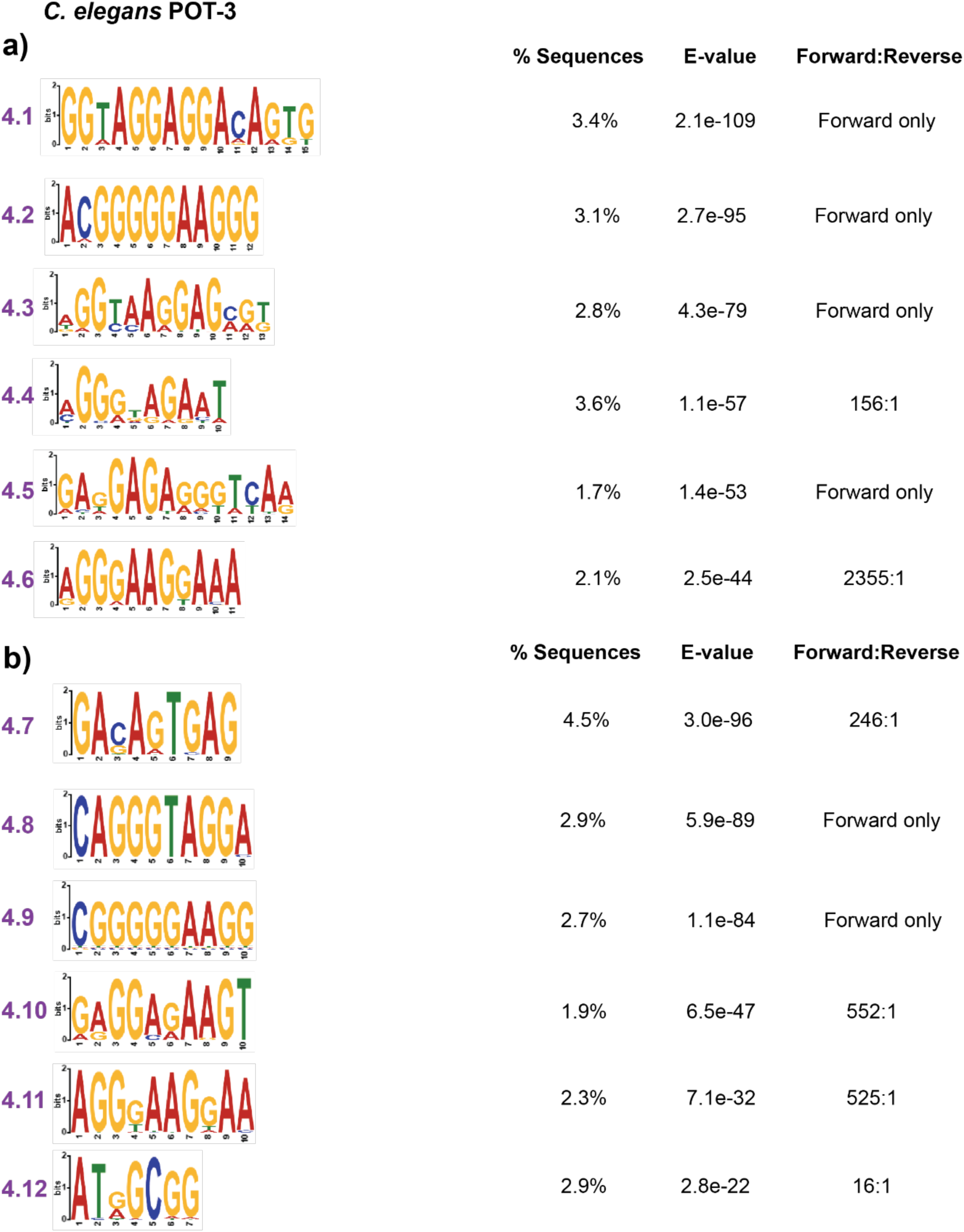
Analysis of HT-SELEX with *C. elegans* POT-3. (a) The six most abundant motifs (listed in decreasing E-value) when searching for motifs 10-25 nts in length. % Sequences indicates the proportion of sequences containing the motif, E-value is an adjusted measure of statistical significance, and Forward:Reverse indicates the relative abundance of the motif in either orientation. See **Methods** section for details regarding these metrics. (b) Same as (a) but showing the results when searching for motifs 5-10 nts in length.

When we searched for motifs 10-25 nts in length in our HT-SELEX dataset generated using 10 ng of MRT-1 for each round of selection, STREME analysis showed that the two most abundant motifs had a total abundance of 33% and enrichment for G nucleotides (**Figure S1a**). The other four most abundant motifs had E-values > 0.05, which called into question their biological relevance for understanding the DNA-binding properties of MRT-1 (data not shown). When examining this dataset for motifs 5-10 nts in length, we found that only four of the most abundant motifs had E-values <0.05: Motifs 5.3-5.6 (**Figure S1b**). Of these, Motifs 5.3, 5.4, and 5.6 show a modest preference for G nucleotides. Given the absence of a full *C. elegans* telomeric repeat among the MRT-1 SELEX consensus sequences^15,16^, it is important to note that MRT-1 contains an SNM1 family nuclease domain^24^. Other members of this family have shown nuclease activity that is stimulated by divalent metal cations and inhibited by EDTA^59^.

Any nuclease activity on the SELEX library DNAs, despite the inclusion of 10 mM EDTA in our SELEX conditions, could confound the interpretation of MRT-1’s DNA binding preference^24,59^. Overall, our study of *C. elegans* proteins using HT-SELEX and our analysis pipeline shows the potential of this procedure to characterize the DNA-binding properties of proteins in an unbiased manner.

## DISCUSSION

We sought to establish a SELEX-based method for determining the DNA-binding properties of telomeric proteins. Having reviewed existing bioinformatics tools, we built a pipeline for SELEX data analysis, beginning with processing of NGS sequencing results (in FASTQ format) and culminating in motif discovery and statistical analysis of enrichment^25,36,37,47^. After performing their SELEX protocol, Choi *et al.* cloned and sequenced fifty of the resulting sequences, demonstrating that hPOT1 enriched the known 10mer ss telomeric DNA binding site^12^ (Class I) as well as a sequence that was partially telomeric and partially non-telomeric (Class II)^5^. This second class suggested that hPOT1 bound to the 5’ phosphorylated dC of a ds-ss telomeric DNA junction, which was confirmed by x-ray crystallography^5,4^.

In the current study, we expanded on this SELEX method by harnessing the high-throughput nature of NGS and bioinformatics analysis to identify motifs in a set of sequences over 1000-fold larger than the dataset analyzed by Choi *et al*^25,36,47,37,5^.

Using this approach for hPOT1, we corroborated the findings of Choi *et al.* by identifying motifs that were akin to Class I and Class II^5^. In so doing, we also validated our *in vitro* and computational approach to characterizing DNA-binding proteins by HT-SELEX. We also explored a new frontier in telomere biology by performing SELEX with *C. elegans* telomeric DNA binding proteins POT-1, POT-2, POT-3, and MRT-1^24,17,20^. Our results suggest that all four *C. elegans* proteins preferentially bind G-enriched sequences. In addition, our analysis of SELEX results for *C. elegans* POT-1 suggests that it binds to DNA hairpins and other secondary structural elements in a manner reminiscent of the way hPOT1 binds the ds-ss telomeric DNA junction^56,4^. Our SELEX data suggest that the OB2-containing *C. elegans* proteins POT-2, POT-3, and MRT-1 exhibit specificity for the G nucleotides within the telomeric repeat rather than showing selectivity for the entire repeat. Intriguingly, our previous analysis of the structural basis of *C. elegans* TEBP-1/-2 binding to telomeric dsDNA did not reveal any protein-based recognition of the “C” base within the G-rich repeat. Thus, both the ds and ss DNA-binding telomeric proteins in *C. elegans* seem to adopt a path of least resistance to conform with the C nucleotide within the G-rich telomeric repeat, by relying on binding to the canonical G nucleotides for sequence specificity. Future structural studies of DNA-bound complexes of *C. elegans* proteins that are considered homologs of human POT1 will allow for a more rigorous testing of this hypothesis.

We present a bioinformatics pipeline that is highly accessible to the biomedical research community. In particular, the tools involved (AptaSUITE^25,36^, STREME^47,37^, Notepad++, R) are freely available and compatible with a Windows OS. In addition, while some simple R code was used in the analysis of STREME results^47^, this R code is available upon request, and the overall pipeline requires little coding experience.

We envision that this pipeline will be applicable *mutatis mutandis* to studies of other telomeric and non-telomeric proteins to determine their nucleic acid-binding properties^7,9,11^. This could include development of aptamers that selectively bind and inhibit proteins that are not canonical DNA/RNA-binding proteins^7^.

Future innovations of this pipeline could include adapting it for the discovery of discontinuous motifs^45,37^. The STREME algorithm is a versatile tool for discovering motifs ranging from three to thirty nucleotides in length; however, it identifies continuous, uninterrupted motifs, which could limit the applications of the algorithm^47,37^. Notably, another resource in the MEME Suite, GLAM2 is suited for finding motifs that include spaces, which could expand the pipeline to characterize more complex and diverse protein-DNA interactions^45,37^. For example, given that POT-1 forms a complex with TEBP-1 and -2^16,22,23^, SELEX analysis of the complex followed by a search for discontinuous motifs could allow for the characterization of the binding site for the entire POT-1/TEBP-1 or POT-1/TEBP-2 complex on telomeric DNA^45,37^.

While we performed secondary structural predictions for a subset of the motifs we discovered in our analysis, this part of the pipeline remains low throughput. We envision that the current pipeline could be adapted to search for candidate secondary structural elements. For example, this approach could begin with searching the SELEX dataset for sequences motifs that are hypothesized to form hairpins (e.g., 5’-GGNNNNCC-3’, 5’-GCNNNNGC-3’, etc.) or that are G-rich. Once a list of candidate sequences is found, we propose grouping them according to sequence similarity using AptaCLUSTER^41,36^. After grouping, each subset would be analyzed using STREME and/or GLAM2^47,45,37^. Finally, consensus sequences derived from the resulting motifs could be entered into a secondary-structure prediction tool, like UnaFold^53,57,58^ or QGRS Mapper^55^. Another exciting modification would be to incorporate secondary-structure prediction for an entire dataset of sequences, ranking each sequence in terms of thermodynamic favorability of potential secondary structures^53,57,58^.

Overall, we have developed and validated an approach for high-throughput analysis of protein-DNA interactions with uses within and beyond the field of telomere biology^60,9,11,7^. We anticipate its further application to understanding the biology of eukaryotes as diverse as humans and nematodes.

## METHODS

### Cloning

*C. elegans* POT-1_N_ comprised amino acids (aa) 1-171, and TEBP-1_PBR_ comprised aa 718-837. For expression of the complex of POT-1_N_ and TEBP-1_PBR_ for SELEX, SUMO-tagged POT-1_N_ and 10xHis-SUMO-tagged TEBP-1_PBR_ were cloned into a pETDuet vector. Full-length POT-2 (aa 1-251) and MRT-1 (aa 1-608) were cloned into a pET28c+ vector to have an N-terminal 6xHis tag. The cDNA for POT-3 was cloned into the pET28b-Smt3 vector to have an N-terminal 10xHis tag.

### Protein Expression and Purification

The DNA-binding domain of human POT1 (hPOT1-DBD) was expressed as a 6xHis-SUMOstar-tagged protein in Hi5 insect cells using a baculovirus-based expression system. The hPOT1-DBD protein was purified as described elsewhere, with modifications described below^4^. Once the protein was eluted from the Ni-NTA agarose resin, it was dialyzed using buffer containing 25 mM Tris pH 8, 500 mM NaCl, and 10 mM 2-mercaptoethanol. Subsequently, protein was adjusted to 10% glycerol and flash frozen.

For co-expression of POT-1_N_ and TEBP-1_PBR_, BL21 (DE3) cells were transformed with the appropriate plasmid, and expression was induced overnight with 100 µM isopropyl β-D-1 thiogalactopyranoside (IPTG) at 25 °C. Cells were sonicated in lysis buffer containing 25 mM Tris pH 8, 500 mM NaCl, 0.1 mM EDTA, 1 mM PMSF, 10 mM 2-mercaptoethanol, and cOmplete mini EDTA-free protease inhibitor cocktail.

Lysate was centrifuged and the resulting supernatant was combined with Ni-NTA agarose beads. Ni-NTA beads were initially washed in Buffer A (25 mM Tris pH 8, 150 mM NaCl, and 10 mM 2-mercaptoethanol), beads were later washed with Buffer A adjusted to 10 mM imidazole and 300 mM NaCl, and protein was eluted with Buffer A adjusted to 300 mM imidazole. The eluted protein was dialyzed in buffer (25 mM Tris pH 8, 150 mM NaCl, and 10 mM 2-mercaptoethanol), adjusted to 10% glycerol, and flash frozen. POT-2 and POT-3 were expressed and purified similarly to the POT-1_N_ and TEBP-1_PBR_ complex. MRT-1 purification included further purification using Superdex 200 size-exclusion chromatography to remove shorter protein contaminants, most likely representing degradation fragments.

### SELEX

Our SELEX procedure was based on that of Choi *et al.*^5^, with the following modifications. The composition of SELEX binding buffer was 25 mM Tris pH 8.0, 100 mM NaCl, 10 mM EDTA, 4% glycerol, 0.01% Triton X-100, and 10 mM 2-mercaptoethanol. (For SELEX with hPOT1-DBD and the POT-1_N_-TEBP-1_PBR_ complex, binding buffer had 1 mM MgCl_2_ instead of 10 mM EDTA.) His-Tag Isolation and Pulldown beads (Dynabeads™, Thermo Scientific) were washed in the appropriate SELEX binding buffer prior to adding them to the protein-DNA mixture (see below). To form protein-DNA complexes in the first round of selection, we combined 5 ug of the N35 library DNA (IDT) with 1 ug of bovine serum albumin (BSA) and the protein of interest in SELEX binding buffer for a total volume of 20 uL. (We used 10 ng of hPOT1-DBD, 10 ng of POT-1_N_-TEBP-1_PBR_ complex, 100 ng of POT-2, 100 ng of POT-3, and either 10 or 100 ng of MRT-1 for each round of selection.) The mixture was incubated on ice for 30 minutes (min), spiked with washed Dynabeads, and nutated at 4 °C for another 30 min. After using SELEX binding buffer to wash the protein-bound beads, ssDNA was eluted by heating 5 min at 95 °C in 1X Phusion HF Buffer. To amplify dsDNA from the eluted ssDNA, some of this elution was combined with a forward primer (5’-CAGTAGCACACGACATCAAG-3’) and reverse primer (5’-CAACTGACACGAGACATGCA-3’) and subjected to PCR^5^. Reaction conditions were optimized by monitoring products by agarose gel electrophoresis. Typical PCR protocols consisted of 30 second (sec) denaturation at 98 °C; 10 sec at 98 °C, 20 sec at 50 °C, and 10 sec at 72 °C, repeated for a total of 9-18 times; and 60-sec extension at 72 °C. To regenerate ssDNA, we conducted asymmetric PCR with forward primer only (16-17 cycles). This ssDNA product was used as input for the next round of selection, after combining with 5 ug of salmon sperm DNA. (Reaction volume for protein-DNA complex formation could be as large as ∼30 uL on subsequent rounds of selection.)

For hPOT1-DBD, we did four rounds of selection, but for all other protein used in SELEX, we did five rounds of selection. Following selection, PCR was performed to add Illumina adaptors and extend oligos to a length of 150 base pairs. After preparing DNAs with KAPA beads (Roche), we subjected them to Amplicon-EZ NGS (Azenta). A sample of unselected N35 library was subjected to the same sequencing protocol.

### SELEX Data Analysis

For the SELEX experiments with human and *C. elegans* proteins, the mean quality scores of the reads were 33 or higher. Paired-end reads were imported in FASTQ format into AptaSuite^36^ and the paired-end reads were merged and sequences of interest extracted using the AptaPLEX tool^25^. Of the ∼460,000 input reads in the hPOT1-DBD dataset, ∼430,000 were accepted, meeting the requirements of sufficient matching to primer sequences and sufficient overlap of paired-end reads. Of the ∼580,000 input reads in the *C. elegans* POT-1_N_-TEBP-1_PBR_ dataset, ∼240,000 were accepted. Of the ∼250,000 reads in the *C. elegans* POT-2 dataset, ∼220,000 were accepted. Of the ∼130,000 reads in the *C. elegans* POT-3 dataset, ∼120,000 were accepted. Of the ∼220,000 reads in the *C. elegans* 10-ng MRT-1 dataset, ∼190,000 were accepted. Of the ∼400,000 reads in the *C. elegans* 100-ng MRT-1 dataset, ∼360,000 were accepted. Extracted sequences for each dataset were exported as a FASTA file and unique numbers were assigned to identify each sequence using Notepad++. The resulting FASTA file for each dataset was used as input to the motif-discovery tool STREME (part of the MEME Suite version 5.5.8) and analyzed using the STREME algorithm on the MEME Suite public web server with motif size range set to either 10-25 nucleotides (nts) or 5-10 nts and the number of motifs discovered capped at 10 or 20 (for human or *C. elegans* proteins, respectively)^37,47^. STREME analyzed a random sample of about 110,000 sequences each for human and *C. elegans* datasets and used Fisher’s exact test for calculations of p-values for motif statistical significance. E-values were calculated by multiplying p-values by the number of motifs identified.

Abundance of each discovered motif (“% Sequences”) was calculated by the STREME algorithm as the percentage of sequences containing at least one instance of the motif. Each time the algorithm iterates over the input sequences, it erases sites where it has already identified motifs. Thus, in cases where motifs overlap and one of the motifs is identified on an earlier iteration of the algorithm, the overlapping site will be included as an instance of the initially identified motif but may not be counted as an instance of a motif discovered in a later iteration. STREME results were further analyzed to determine orientation of the motifs in the input dataset using R. Briefly, output of the STREME algorithm (a tsv file containing each occurrence of the motif in the analyzed sequences) was used as input to an R function that counted and graphed the number of occurrences that a motif appeared in forward orientation on the ssDNA or as its reverse complement in the sequences. (Instances of each motif that were tabulated in the tsv files were counted in a way that would count sites where motifs overlapped as examples of each of the overlapping motifs. Thus, unlike the % Sequences calculation, these motif instances were determined without site erasure.) The R code of this function is available upon request. For all figures that display sequence logos, the logo for the orientation that predominated was chosen. The ratios of forward vs. reverse orientation were calculated by dividing the larger count by the smaller count and rounding to the nearest whole number. Predictions of G-quadruplex formation were made using the QGRS Mapper tool^55^. Predictions of DNA secondary structure was performed using _UNAFold53,57,58._

## ACKNOWLEDGMENTS.

We thank T. L. Bailey (University of Nevada at Reno, USA) and C. E. Grant (the MEME Suite) for information related to the MEME Suite. We thank J. Hoinka (National Library of Medicine, National Institutes of Health) for information related to AptaSUITE. This work was supported by National Institutes of Health grant R35GM148276 (J.N.) and National Science Foundation grant project 2425568 (J.N.). J.W. was supported by NIH T32 GM145470.

