## Supplemental Figure 1 for "A Data-Analysis Pipeline for High-Throughput Systematic Evolution of Ligands by Exponential Enrichment (HT-SELEX) in the Characterization of Telomeric Proteins"

**Supplementary Figure legend**

**Figure 1.** Analysis of HT-SELEX with *C. elegans* MRT-1. (a) The two most abundant motifs (listed in decreasing E-value) when searching for motifs 10-25 nts in length. % Sequences indicates the proportion of sequences containing the motif, E-value is an adjusted measure of statistical significance, and Forward:Reverse indicates the relative abundance of the motif in either orientation. See **Methods** section for details regarding these metrics. (b) Same as (a) but showing the four most abundant motifs when searching for motifs 5-10 nts in length. We also performed HT-SELEX using 100 ng of MRT-1 per round of selection (data not shown). When searching this dataset for motifs 5-10 nts in length, we saw a motif containing 5'-AGGGTT-3' (a permutation of the mammalian telomeric repeat[1]) with an abundance of ~3%. Otherwise, our analysis yielded similar results as for the 10-ng dataset.

### Figure S1

*C. elegans* MRT-1

**a)**

| Motif | Logo | % Sequences | E-value | Forward:Reverse |
|---|---|---|---|---|
| 5.1 | 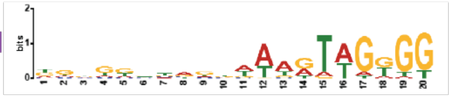 | 26.0% | 1.1e-5 | 13:1 |
| 5.2 | 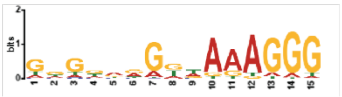 | 7.0% | 2.5e-5 | 33:1 |

**b)**

| Motif | Logo | % Sequences | E-value | Forward:Reverse |
|---|---|---|---|---|
| 5.3 | 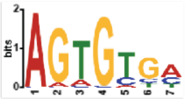 | 8.1% | 1.3e-3 | 4:1 |
| 5.4 | 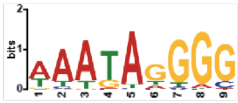 | 7.5% | 3.2e-3 | 8:1 |
| 5.5 | 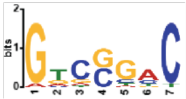 | 5.0% | 3.2e-2 | N/A (palindrome) |
| 5.6 | 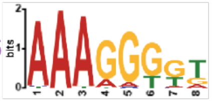 | 4.3% | 4.3e-2 | 14:1 |
